# Senolytic-Resistant Senescent Cells Have a Distinct SASP Profile and Functional Impact: The Path to Developing Senosensitizers

**DOI:** 10.1101/2025.08.27.672709

**Authors:** Utkarsh Tripathi, Masayoshi Suda, Vagisha Kulshreshtha, Bryan T. Piatkowski, Allyson K. Palmer, Nino Giorgadze, Christina Inman, Nathan Gasek, Ming Xu, Kurt O. Johnson, Tamar Pirtskhalava, Selim Chaib, Larissa P. G. Langhi Prata, Yi Zhu, Renuka Kandhaya-Pillai, Stefan G. Tullius, Saranya P Wyles, Rambabu Majji, Hari Krishna Yalamanchili, David B. Allison, Tamar Tchkonia, James L. Kirkland

## Abstract

The senescent cell (SC) fate is linked to aging, multiple disorders and diseases, and physical dysfunction. Senolytics, agents that selectively eliminate 30-70% of SCs, act by transiently disabling the senescent cell anti-apoptotic pathways (SCAPs), which defend those SCs that are pro-apoptotic and pro-inflammatory from their own senescence-associated secretory phenotype (SASP). Consistent with this, a JAK/STAT inhibitor, Ruxolitinib, which attenuates the pro-inflammatory SASP of senescent human preadipocytes, caused them to become “senolytic-resistant”. Administering senolytics to obese mice selectively decreased abundance of the subset of SCs that is pro-inflammatory. In cell cultures, the 30-70% of human senescent preadipocytes or human umbilical vein endothelial cells (HUVECs) that are senolytic-resistant (to Dasatinib or Quercetin, respectively) had increased p16^INK4a^, p21^CIP1^, senescence-associated β-galactosidase (SAβgal), γH2AX, and proliferative arrest similarly to the total SC population (comprising senolytic-sensitive <u>plus</u> -resistant SCs). However, the SASP of senolytic-resistant SCs entailed less pro-inflammatory/ apoptotic factor production, induced less inflammation in non-senescent cells, and was equivalent or richer in growth/ fibrotic factors. Senolytic-resistant SCs released less mitochondrial DNA (mtDNA) and more highly expressed the anti-inflammatory immune evasion signal, glycoprotein non-melanoma-B (GPNMB). Transplanting senolytic-resistant SCs intraperitoneally into younger mice caused less physical dysfunction than transplanting the total SC population. Because Ruxolitinib attenuates SC release of pro-apoptotic SASP factors, while pathogen-associated molecular pattern factors (PAMPs) can amplify the release of these factors rapidly (acting as “senosensitizers”), senolytic-resistant and senolytic-sensitive SCs appear to be interconvertible.

## Introduction

The senescent cell (SC) fate is linked to aging, multiple disorders and diseases, and physical dysfunction [1-9]. Senescence can occur in most cell types across the vertebrates and can be induced by replicative stress, DNA damage, cytotoxic drugs, radiation, inflammation, metabolic dysfunction, pathogen exposure, and other insults [1-3, 5, 6, 9]. Senescence entails resistance to apoptosis and SCs are generally removed by innate and adaptive immune system components [10, 11]. Senescence-associated cell cycle arrest can serve to curtail the proliferation of dysfunctional, damaged, or pre-cancerous cells [12]. In preclinical cell and tissue culture and animal models, it appears that while senescence contributes to a range of pathological processes, this cell fate has essential roles, such as for cancer prevention, embryonic development and parturition, and wound healing [4, 10, 13-16]. However, those persisting SCs that are not removed by the immune system can develop a tissue-destructive SASP, spread senescence locally and systemically, develop and harbor potentially cancerous mutations, and can lead to deleterious sequelae, including tumor development, chronic inflammation, immune deficits, and progenitor cell dysfunction [15, 17-21]. Consistent with this, transplanting small numbers of SCs can result in frailty, osteoarthritis, and accelerated death from most of the diseases that occur later in life in naturally aged mice [22, 23].

SCs are not homogeneous: they exhibit significant heterogeneity in their characteristics and behavior depending on the type of cell that became senescent, the inducer of senescence, time since senescence was induced, host *milieu*, and disease states and arguably, SCs may be of pro-inflammatory “deleterious” or reparative “helper” subtypes [24-27]. There are challenges in examining the heterogeneity of SCs across human tissues due to the low abundance of SCs and controversy about sensitive and specific markers of senescence [28].

Senolytics are agents that were originally developed using a hypothesis-driven, mechanism-based approach to target and eliminate senescent cells selectively: the first senolytics were designed to transiently disable the SCAPs that protect the subset of SCs that are pro-apoptotic from being targeted and removed by their own SASP [29-31]. We previously demonstrated that the first senolytics reported, Dasatinib and Quercetin, eliminate 30-70% of senescent cells through apoptosis (terminal deoxynucleotidyl transferase dUTP nick end labeling [TUNEL] assay; Figs. 2E&F in [29]) and percent of cleaved caspase 3^+^ cells (Fig. 4E in [32]). Although Dasatinib and Quercetin remove 30-70% of SCs, senolytics still delay, prevent, alleviate, or treat multiple disorders and diseases across the lifespan in pre-clinical models [22, 29, 33-40]. However, there are few data about the nature and role of the “senolytic-resistant” SCs that remain after senolytic exposure *vs*. “senolytic-sensitive” SCs. Here, we explore mechanisms through which those SCs that are sensitive *vs*. resistant to senolytics can be interconverted and their impact *in vivo* by transplanting these two senescent cell subtypes into younger mice.

## Materials and Methods

### Cell Culture

Preadipocytes (also known as adipose-derived stromal cells or adipose mesenchymal “stem” cells [MSCs]) were isolated from abdominal subcutaneous fat biopsies obtained from subjects donating kidneys for transplantation aged 38.4 +/- 11.4 years (mean +/- SD), median 37 (25–68), with a body mass index 28.3 +/- 3.9 (mean +/- SD), median 27.2 (23-40.46) or undergoing bariatric surgery aged 39.25 +/- 8.5 years (mean +/- SD), median 40.5 (28-48) with a body mass index 39.7 +/- 5.03 (mean +/- SD), median 37.29 (36.6-47.1), both males and females. All subjects gave informed consent. The protocol was approved by the Mayo Clinic Institutional Review Board for Human Research. Using methods we previously reported [27]: 1) human preadipocytes were isolated and cultured; 2) all human preadipocyte lots that were not induced to become senescent retained proliferative capacity over multiple passages with low markers of senescence, and were therefore considered to be non-senescent/ control cells; and 3) human preadipocyte senescence was induced by 20 Gy X-irradiation and experiments were performed 30 days after radiation. To minimize inter-donor variability, we employed a paired experimental design in Figures 1B, D, 4E, 5B, S4C, and S4D. Preadipocytes from each donor were divided into three groups: one served as a non-irradiated control (non-senescent) and the others were subjected to irradiation to induce senescence followed by vehicle or senolytic treatment. In HUVECs purchased from Lonza (CC-2519, San Diego, California), senescence was induced by 10 Gy X-irradiation and experiments were conducted 10-14 days after irradiation. The MDA-MB-231 human breast cancer cell line was obtained from ATCC. Cells were treated with 1µ/ml Cisplatin for 24 hrs and maintained in culture for 1 week. The cells were then treated with Dasatinib (200nM) or vehicle for 24 hrs followed 24 hrs later by a TLR3 agonist, polyinosine-polycytidylic acid (10 µg/ml), or vehicle daily for 3 days. After treatment with the TLR3 agonist or vehicle, cells were again exposed to Dasatinib or vehicle for 24 hrs.

**Fig. 1.**
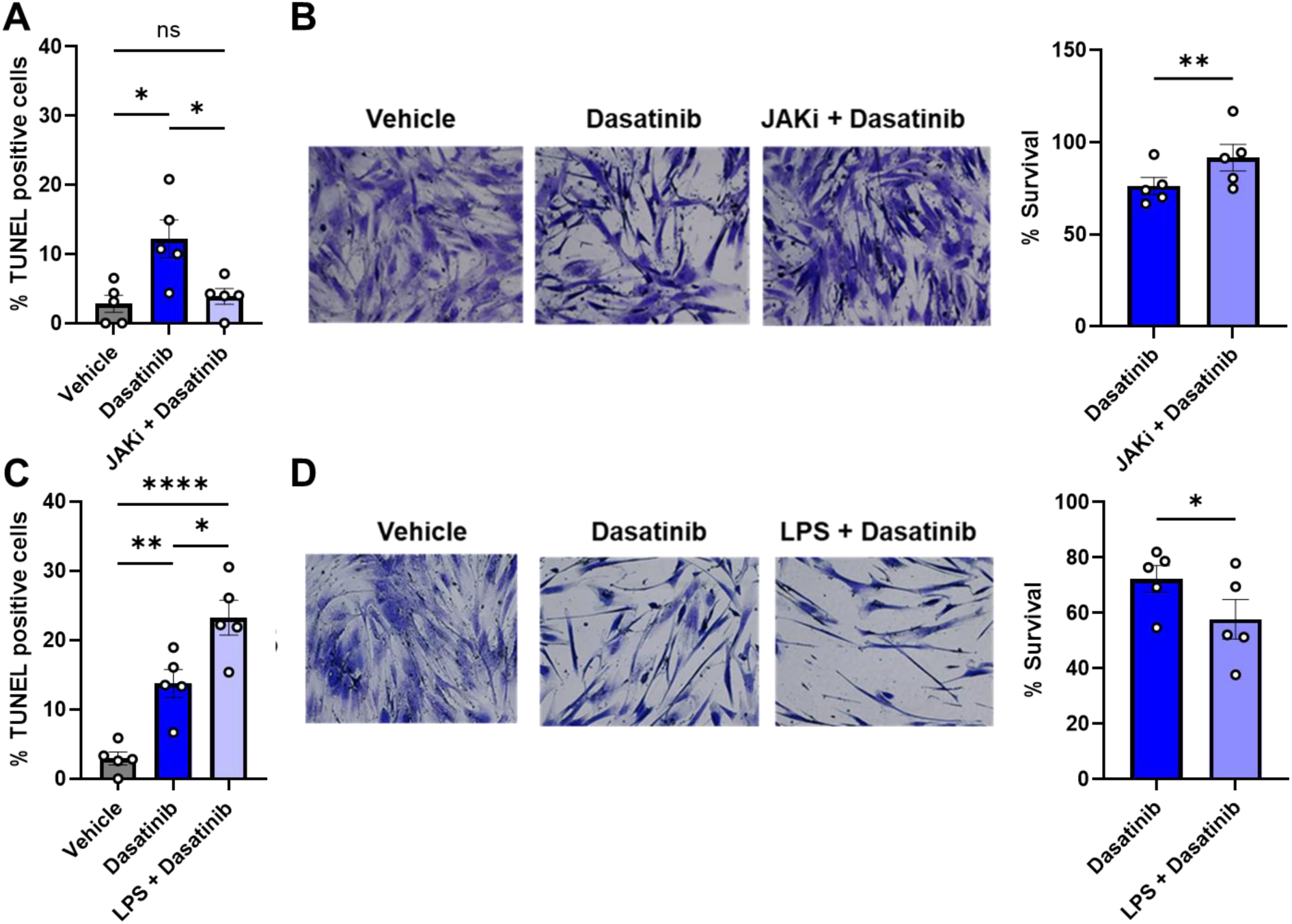
The SASP impacts extent of SC clearance by senolytics. (A) TUNEL-positive nuclei as a percent of total cells and (B) representative images of surviving (crystal violet^+^) senescent preadipocytes and quantification relative to vehicle-treated cells after exposure to vehicle or Ruxolitinib (1µM), which attenuates the pro-inflammatory SASP, for 3 days followed by Dasatinib 800 nM for 24 hrs. (C) TUNEL-positive nuclei as a percent of total cells and (D) survival of SCs pre-treated with 10 µg/ml LPS for 3 days followed by treatment with Dasatinib 800 nM for 24 hrs. Quantification of images (N=5) is shown on the right. Data are shown as means +/- SEM; 1-way ANOVA; and *post hoc* comparisons with by Tukey’s HSD multiple comparison (A, C), and are expressed as a function of vehicle-treated cells; means +/- SEM; paired, 2-tailed Student’s T-tests.

### Reagents

Dasatinib (SML2589) and Quercetin (Q4951) were purchased from Sigma (Sigma, St. Louis, MO, USA). JAK inhibitor, Ruxolitinib (tlrl-rux) was purchased from InvivoGen (Sigma, St. Louis, MO, USA). Lipopolysaccharide (LPS; tlrl-3pelps) was purchased from InvivoGen (San Diego, CA, USA).

### rt-PCR and RNA Sequencing

For most studies, cells were washed with PBS and RNA was isolated using Trizol and chloroform [41]. Concentration and purity of samples were assayed using a Nanodrop spectrophotometer. Each cDNA sample was generated by reverse transcription using 1-2,000 ng RNA following the manufacturer’s recommended protocol (High-capacity cDNA Reverse Transcription Kit; Cat #4368813, Thermo Fisher Scientific, Waltham, MA, USA). A reverse transcription program was used (10 min. at 25°C, 120 min. at 37°C, 5 min. at 85°C, held at 4°C). TATA-box binding protein (TBP) served as a control for gene expression analyses. A list of primers is provided in Table 2. For RNA sequencing, library preparation was performed using a TruSeq Stranded mRNA kit prior to sequencing on the Illumina NovaSeq 6000 platform.

Raw reads were trimmed with fastp (v0.20.1) [42]. Trimmed reads were then mapped to the reference genome GRCh38 with STAR (v2.7.9a) [43] and gene expression is estimated by generating a count matrix with STAR-counts. The raw genes counts were adjusted for the sample pairing covariate using ComBat-Seq within the R package, sva v3.50.0 [44]. Differential expression analysis was performed using the R package, edgeR v4.0.16 [45]. Raw gene counts were first pre-filtered to keep only those genes with greater than one average count per million in either comparison group. The raw *p*-values were adjusted using the Benjamini-Hochberg procedure [46] to control the false discovery rate and those genes with an adjusted *p*-value ≤ 0.05 were considered to be significant. Differentially expressed genes were used for heatmap visualization and hierarchical clustering. Functional enrichment analyses were performed using clusterProfiler v4.14.0 [47].

### Conditioned Media (CM)

CM were prepared by exposing cells to RMPI 1640 containing 1 mM sodium pyruvate, 2 mM glutamine, MEM (minimum essential medium) vitamins, MEM nonessential amino acids (all Gibco), and Streptomycin (Life Technologies, Carlsbad, CA, USA) [41]. CM were collected 24-48 hrs. after the cells had been exposed to the media.

### TUNEL Assay

Cellular apoptosis was assessed using the In Situ Cell Death Detection Kit (Roche, 11684795910) according to the manufacturer’s instructions. Samples were counterstained with DAPI (Life Technologies). Apoptotic and non-apoptotic cells were examined by fluorescence microscopy (Nikon ECLIPSE Ti, NIS Elements AR 5,20,02).

### Cell Survival

Cell survival was assayed by crystal violet. Briefly, cells were washed with PBS twice and fixed with 4 % PFA in PBS for 15 mins. and then washed again with PBS after incubation. Cells were then treated with 0.5% crystal violet in 25% methanol. After incubation for 10 min., crystal violet was removed and plates were washed with water until washings were clear of visible dye. Plates were dried and crystal violet was dissolved in 100% methanol. Absorbance at 570 nM was quantified.

### SAβgal Assay

SAβgal activity was assayed as previously described [48]. In brief, primary preadipocytes were washed with PBS and then fixed for 5 mins. in PBS containing 2% (vol/vol) formaldehyde (Sigma-Aldrich, ST. Louis, USA) and 0.25% glutaraldehyde (Sigma-Aldrich, St. Louis, USA). Following fixation, cells or tissues were washed with PBS before being incubated in SAβgal activity solution (pH 6.0) at 37°C for 16-18 hrs. The enzymatic reaction was stopped by washing cells or tissues 3-5X with ice-cold PBS. Bright field microscopy (Nikon Eclipse Ti) was used for imaging.

### mtDNA Assay

Total DNA was extracted from media using the QIAmp-Blood & Tissue kit (Qiagen, Germantown, MD, USA). mtDNA copy number was measured by quantitative rtPCR using primers for mitochondrially encoded NADH dehydrogenase subunit 2 (MT-ND2) and cytochrome c oxidase subunit III (MT-COX3) of the electron transport chain.

### Immunostaining

Cells were fixed with 4% (vol/vol) paraformaldehyde for 10 mins. and permeabilized using PBS containing 0.25% Triton X-100 for 10 mins. After being washed with PBS, cells were blocked with PBS and Tween-20 (PBST) containing 1% BSA for 30 mins. Cells were incubated with primary antibodies in 1% BSA in PBST (blocking buffer) overnight at 4°C, washed with PBS and then incubated with secondary fluorescent antibodies (Life Technologies, Carlsbad, CA, USA) in the blocking buffer for 1 hr. in the dark. DAPI (Life Technologies, Carlsbad, CA, USA) was used to stain nuclei for cell counting. Fluorescence microscopy (Nikon Eclipse Ti) was used for imaging. For EdU incorporation assays, cells were incubated in 10 μM EdU for 24 hrs. in growth medium. All subsequent EdU detection steps were carried out using Click-iT EdU Alexa Fluor Imaging Kits (C10337, Thermo Fisher Scientific, Waltham, MA USA) according to the manufacturer’s instructions. The following antibodies were used: p16 (1:200 dilution, Cat# 705-7493, Cintech Histology); p21 (1:100 dilution, Cat# 109199, Abcam, Cambridge, MA); and Phospho-Histone H2A.X (Ser139) (20E3) (Rabbit; 1:200 dilution, Cat #9718, Cell Signaling, Danvers, MA).

### Mice

Two-month-old *SCID-beige* mice (Charles River Laboratories, Wilmington, MA, USA) were used for the SC transplantation studies. Mice were maintained in a pathogen-free facility at 23-24°C under a 12-hrs. light, 12 hrs. dark regimen with free access to water and a chow diet (standard mouse diet with 20% protein, 5% fat [13.2% fat by calories], and 6% fiber; Lab Diet 5053, St. Louis, MO). For the obese mouse studies, animals were maintained on a 60% (by calories) fat diet (D12492, irradiated; Research Diets, New Brunswick, NJ). All mouse studies were approved by the Mayo Clinic Institutional Animal Care and Use Committee.

### Cell Transplantation

Human preadipocyte senescence was induced by 20 Gy x-irradiation. SCs or control non-irradiated cells were collected by trypsinization. Cell pellets were washed with PBS once and re-suspended in PBS for transplantation. Recipient *SCID-beige* mice were anesthetized using isoflurane and cells were injected intraperitoneally in 150-200 μl PBS through a 22 G needle [22].

### Physical Function Assays

Forelimb grip strength was determined in the transplanted mice using a grip strength meter (Columbus Instruments, Columbus, Ohio) [22]. Results were averaged from 3-5 trials. For the wire hanging test [22], mice were placed on a 2 mm thick metal wire, which was placed 35 cm above a padded surface. Mice were allowed to grab the wire only with their forelimbs. Hanging time was normalized to body weight as hanging duration in seconds × body weight (grams). Results were averaged from 3 trials for each mouse. In this experiment, mice were housed in cages. This may introduce clustering effects due to the dependency of the mice in the same group. The authors acknowledge the need to properly account for clustering due to the risk of underestimating standard errors and inflating the Type I error rates. However, we do not expect a significant impact of the clustering effects in this study because animals from all three groups (treated with non-senescent cells, senescent cells, and senolytic-resistant senescent cells) were together in the same cages. Thus, any within-cluster effects on the variability of the outcome would impact all mice rather than a specific mouse group. Additionally, findings rely more on treatment group responses, not individual mouse responses.

### Multiplex ELISA

CM were filtered and cytokine and chemokine protein levels in CM were assayed using Luminex xMAP technology. Multiplexing analysis was performed using a Luminex 100 system (Luminex, Austin, TX, USA) by Eve Technologies Corp. (Calgary, Alberta, Canada). Data are represented as pg/ml for each SASP factor as a function of cellular density.

### Mass Cytometry by Time of Flight (CyTOF)

An antibody panel was designed based on surface markers, transcription factors, and cytokines. Each antibody was tagged with a rare metal isotope (Table 1) and function verified by mass cytometry according to the manufacturer’s manual (Multi Metal labeling kits; Fluidigm South San Francisco, CA, USA). A CyTOF-2 mass cytometer (Fluidigm, South San Francisco, CA, USA) was used for data acquisition. Acquired data were normalized based on normalization beads (Ce140, Eu151, Eu153, Ho165, and Lu175 as in [49]). Preadipocytes were isolated from mouse epididymal white adipose tissue. Collected cells were incubated with metal-conjugated antibodies for testing intracellular proteins including transcription factors and cytokines. Fixation and permeabilization were conducted according to the manufacturer’s instructions (Foxp3/Transcription Factor Staining Buffer Set, eBioscience, San Diego, CA, USA). CyTOF data were analyzed by Cytobank (Santa Clara, CA, USA). Total preadipocytes were considered to be CD45^−^, CD31^−^, and Sca1^+^, macrophages F4/80^+^ and CD11b^+^, endothelial cells CD31^+^ and CD146^+^, and T lymphocytes CD4^+^ or CD8^+^ as in [49].

**Table 1.** Antibodies used for CyTOF.

| Label | Protein | Clone | Company |
| --- | --- | --- | --- |
| 143 Nd | pS139 H2AX | N1-431 | BD Bioscience <sup>249</sup> |
| 144 Nd | p21 | F-5 | Santa Cruz <sup>250</sup> |
| 145 Nd | CENP-B | ab25734 | Abcam <sup>251</sup> |
| 150 Nd | pS63 c-Jun | 9261 | Cell Signalling <sup>252</sup> |
| 152 Sm | CD3e | 145-2C11 | Fluidigm <sup>253</sup> |
| 153 Eu | CD8a | 53-6.7 | Fluidigm <sup>254</sup> |
| 154 Sm | CD11b | M1/70 | Fluidigm <sup>255</sup> |
| 155 Gd | IL-6 | MP5-20F3 | Biolegend <sup>256</sup> |
| 156 Gd | Activin A | 69403 | R&D Systems <sup>257</sup> |
| 158 Gd | PY705Stat3 | 4/P-Stat3 | Fluidigm <sup>258</sup> |
| 166 Er | CXCL1 | MAB453 | R & D Systems <sup>259</sup> |
| 170 Er | Flag | L5 | Biolegend <sup>260</sup> |
| 171 Yb | IL10 | JES5-16E3 | Biolegend <sup>261</sup> |
| 175 Lu | pS536 NF-κB | 93H1 | Cell Signalling <sup>262</sup> |
| 176 Yb | TNFα | MP6-XT22 | Biolegend <sup>263</sup> |

**Table 2.**
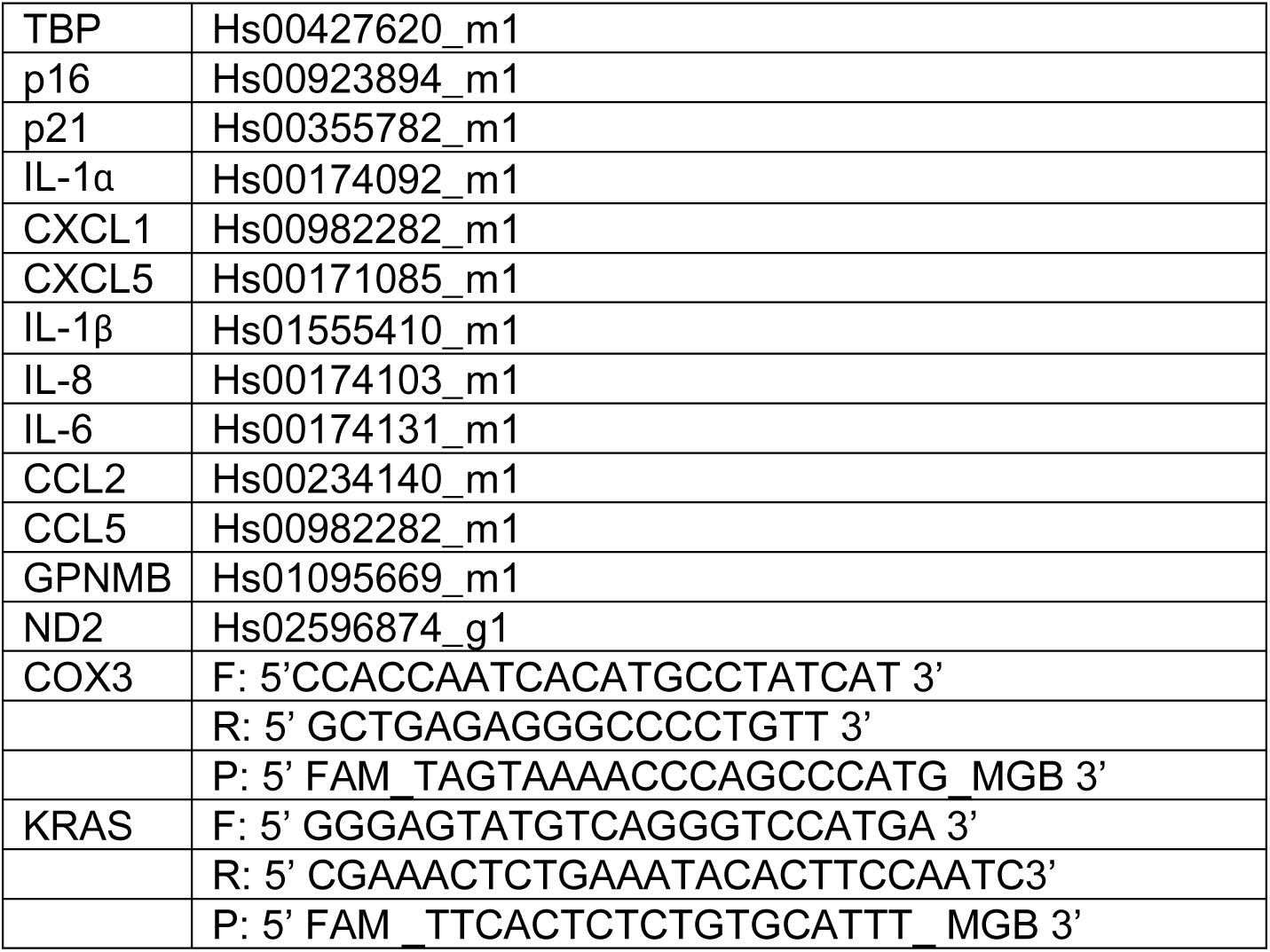
Primers used for rt-PCR analyses.

## Statistical Analyses

GraphPad Prism version 10.0.3 was used for statistical analyses. Results are presented as mean ± SD or SEM as stated in the figure legends. p<0.05 was considered to be statistically significant and p<0.05, p<0.01, p<0.005, and p<0.001 are indicated as *, **, ***, and ****, respectively. Statistical analyses were performed using two-tailed tests. Statistical tests include Student’s T-tests or two-tailed Mann-Whitney tests comparing two groups, one-way ANOVA and *post hoc* multiple comparison test with Tukey’s Honestly Significant Difference (HSD) comparing more than two groups of pairs as detailed in the figure legends. Due to the limited sample size and the dependence of the methods used on the distributional assumptions of normality, a sensitivity analysis was then conducted using nonparametric tests based on taking the ranks of the variables then analyzing the ranks. Statistical significance of the results was diminished upon non-parametric testing as opposed to using standard T tests, supporting the desirability of conducting future confirmatory studies with larger sample sizes.

## Results

### The SASP is Linked to Extent of SC Clearance by Senolytics

The JAK/STAT inhibitor (JAKi), Ruxolitinib, attenuates the SASP of senescent human preadipocytes [41]. Reducing the SASP by Ruxolitinib or si-RNA mediated knockdown of JAK1 attenuated the ability of Dasatinib to clear senescent human preadipocytes (Figs. 1A, B S1A, B). Pathogen-associated molecular pattern (PAMP) factors, such as lipopolysaccharide (LPS), which is in the outer membrane of gram-negative bacteria, has been shown to exacerbate pro-inflammatory SASP factor expression (Fig. S1C) [50]. Here, we found that LPS pre-treatment of SCs enhanced their killing by senolytics (Fig. 1C, D). Hence, interference with SASP factor expression related to the JAK/STAT pathway decreases the impact of senolytic treatment on human preadipocytes, while “senosensitizing” microenvironmental factors such as those linked to infections increase susceptibility of senolytic-resistant SCs to senolytic treatment [27, 50].

### Senolytics Target the Pro-Inflammatory/ Pro-Apoptotic Subset of Senescent Preadipocytes in Obese Mice

Senolytics were developed based on the observation that those SCs that are tissue-damaging employ anti-apoptotic, pro-survival pathways (SCAPs) to protect themselves from their own pro-apoptotic SASP [29]. Different senolytics discovered using this hypothesis-driven approach target 30 to 70% of senescent cells *in vitro* [29, 31]. Here we tested if it is indeed the senescent cells with high pro-inflammatory/ pro-apoptotic SASP factor expression that are selectively vulnerable to senolytics *in vivo*. Fourteen-to 16-month-old male mice that had been on a high-fat diet (see Methods) for 8 months were administered Dasatinib (5 mg/kg) plus Quercetin (50 mg/kg) or vehicle (60% Phosal, 10% ethanol, and 30% PEG-400) for five consecutive days by oral gavage. Mice were euthanized 3 days after the final dose. Organs were harvested and single-cell suspensions from epididymal fat tissue were analyzed by CyTOF (Fig. 2A). Changes in cellular composition caused by senolytics are shown in Fig. 2B. Senolytics decreased preadipocyte and immune cell abundance (black arrows). Markers of senescence, including p16^Ink4a^, p21^Cip1^, CENP-B, and γ-H2AX, were detected in CD34^+^ cells (preadipocytes) and F4/80^+^ cells (macrophages; Fig. 2C). Among preadipocytes, cluster c, a pro-inflammatory/ pro-apoptotic subset of senescent cells, was decreased by senolytics (Figs. 2D, E). With the sample size used in our study, senolytic treatment may have had a slight but yet non-significant effect on macrophages, including those that express senescence markers and pro-inflammatory factors (Figs. S2A, B, S3A). Natural killer (NK) cells were reduced (Fig. S3A), but these NK cells did not have detectible expression of the senescence marker, γ-H2AX (Fig. 2C). Senolytic treatment did not lead to statistically significant changes in endothelial cell abundance, which may relate to their lower expression of senescence markers (Fig. 2D). As we reported previously in cultured cells and tissue explants [29, 32], these findings suggest senolytics primarily target those SCs with a pro-inflammatory/ apoptotic SASP *in vivo* and that presence of pro-inflammatory SCs correlates with immune cell infiltration, however this needs further investigation.

**Fig. 2.**
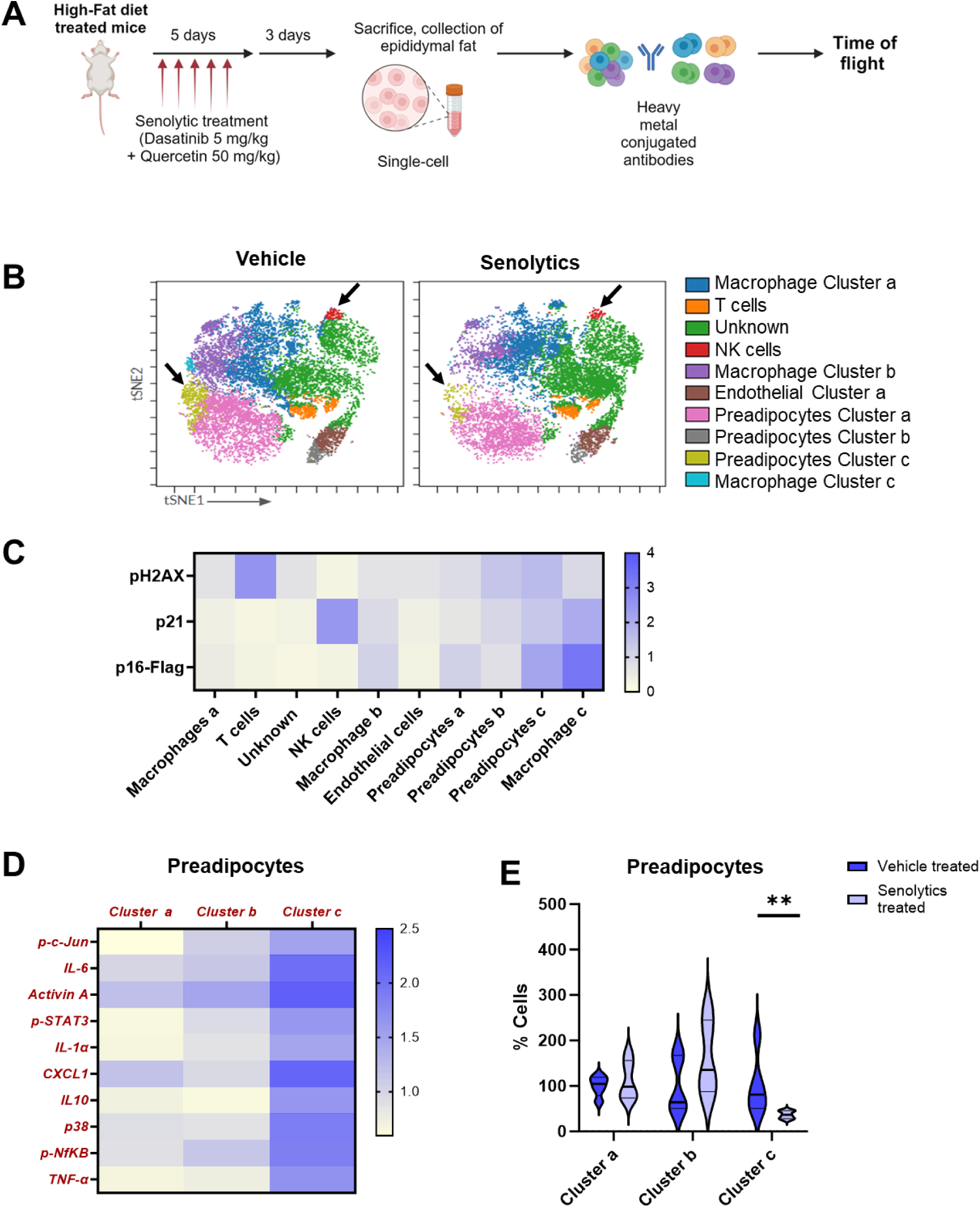
Senolytics target a pro-inflammatory subset of senescent preadipocytes in obese mice. (A) CyTOF experimental scheme. (B) t-SNE plots of FlowSOM clusters of cells from obese vehicle- or senolytic-treated mice (N=5). (C) Heatmap of senescence marker expression within FlowSOM clusters. (D) Heatmap comparing expression of pro-inflammatory SASP factors among preadipocyte clusters in vehicle-treated obese mice. (E) Percent of cells cleared in the indicated clusters in senolytic-*vs*. vehicle-treated mice. Means +/- SEM; unpaired two-tailed Mann-Whitney tests.

### Markers of Cellular Senescence Do Not Differ Significantly Between the Senolytic-Resistant and Total SC Populations

To test whether those cells that are resistant to senolytics are indeed senescent, several established markers of senescence were analyzed. We previously showed that the efficacy of different senolytics depends on the cell type of origin [29]. For example, Dasatinib targets senescent preadipocytes, while Quercetin targets senescent HUVECs [29]. Senescent preadipocytes and HUVECs were treated with Dasatinib or Quercetin, respectively, for 24 hrs. Next, the remaining cells were washed with PBS 4X and subsequently incubated for 3 days. The resulting senolytic-resistant SCs were proliferatively arrested as confirmed by BrdU labeling (Fig. 3A). Senolytic-resistant (only cells remaining after exposure to senolytics) were compared to total (senolytic-resistant *plus* senolytic-sensitive) SC populations, because it is not yet feasible *a priori* to isolate only senolytic-sensitive SCs. In both the total and senolytic-resistant SC populations, we did not detect statistically significant differences in the markers associated with cell cycle inhibition (p16^INK4a^ and p21^CIP1^) and the DNA damage marker, γ-H2AX (Figs. 3B, C, D). Like the total SC population, senolytic-resistant SCs were SAβgal+ (Fig. 3E). Furthermore, senescence markers in Quercetin-treated senescent endothelial cells (HUVECs) were similar to the total senescent HUVEC population (Fig. S4). Hence, consistent with our previous reports [29, 32], we propose, at least *in vitro*, that SC populations comprise two subsets, the one being susceptible to senolytics and the other resistant.

**Fig. 3.**
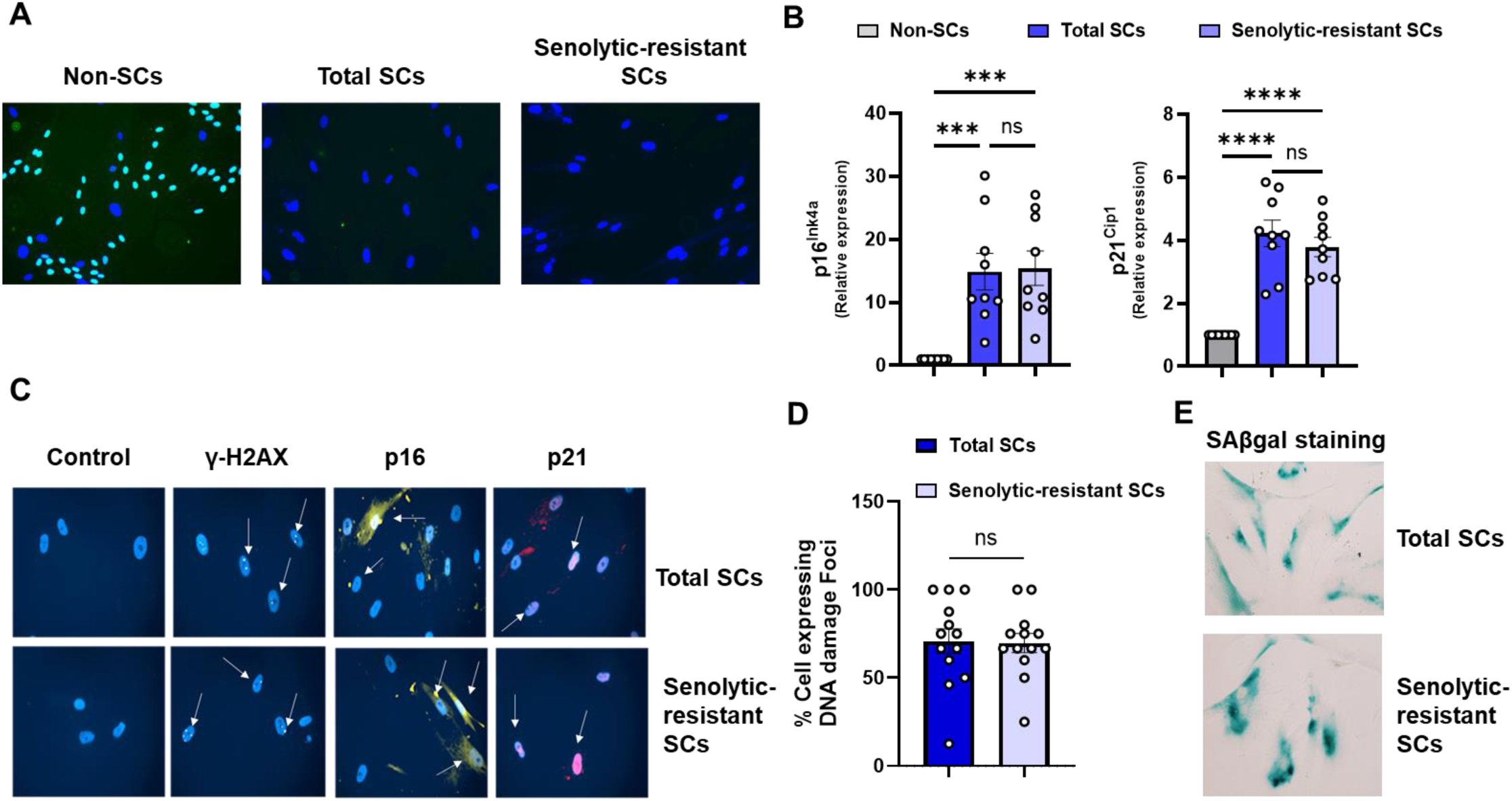
Cellular senescence markers are similar in the human preadipocyte senolytic-resistant *vs*. total SC preadipocyte populations. (A) Cells were analyzed for proliferative arrest by BrdU staining. (B) Gene expression for the indicated senescence markers (N=9) in total senescent preadipocyte populations *vs*. senolytic-resistant senescent preadipocytes. Data are expressed as a function of non-senescent control cells. Means +/- SEM; unpaired, 1-way ANOVA; *post hoc* comparisons by Tukey’s HSD multiple comparison test. (C) Total and senolytic-resistant preadipocyte SC populations were immunostained for γ-H2AX, p16^INK4a^, and p21^CIP1^; a representative image of N=3 subjects is shown. (D) Quantification of γ-H2AX expression. Means +/- SEM; unpaired 2-tailed Student’s T-tests. (E) SAβgal (pH6) in the total and senolytic-resistant senescent preadipocyte populations. Analogous findings in human senescent endothelial populations are in Fig. S3.

### The SASP of the Senolytic-Resistant Is Distinct from That of the Total SC Population

To test further if senolytic treatment indeed targets pro-inflammatory senescent subtypes, bulk RNA-sequencing was conducted of preadipocytes cultured from six healthy human kidney transplant donors after inducing senescence by X-radiation. Clustering of the differential gene expression profiles between senolytic-resistant *vs*. total SCs is shown in the heat map and volcano plots of key genes (Figs. 4A, B). A detailed list of all differentially expressed genes is provided in supplemental Table 1 Potential differences in biological function between the senolytic-resistant *vs*. total human preadipocyte SC populations were ascertained by Gene Ontology over-representation analysis of the differentially expressed genes (Figs. 4C, D, supplemental Table 2). Senolytic-resistant SCs exhibited enrichment of biological processes associated with growth and repair and tissue development compared to the total SC population (Figs. 4D). Repressed functions in senolytic-resistant SCs included activation and migration of immune cells, as further suggested by gene expression and protein analyses (Figs. 4C, E, S5), including genes encoding chemokines/ cytokines such as CXCL1, CXCL5, and CXCL8. Furthermore, senolytic-resistant SCs had higher glycoprotein non-melanoma-B (GPNMB) expression, a recently identified senescence marker [51, 52], compared to the total SC population (Fig. 4E). GPNMB can impact innate and adaptive immune function, decrease inflammation, promote tumor growth, and shield some cancers from immune/ inflammatory defenses [53, 54].

**Fig. 4.**
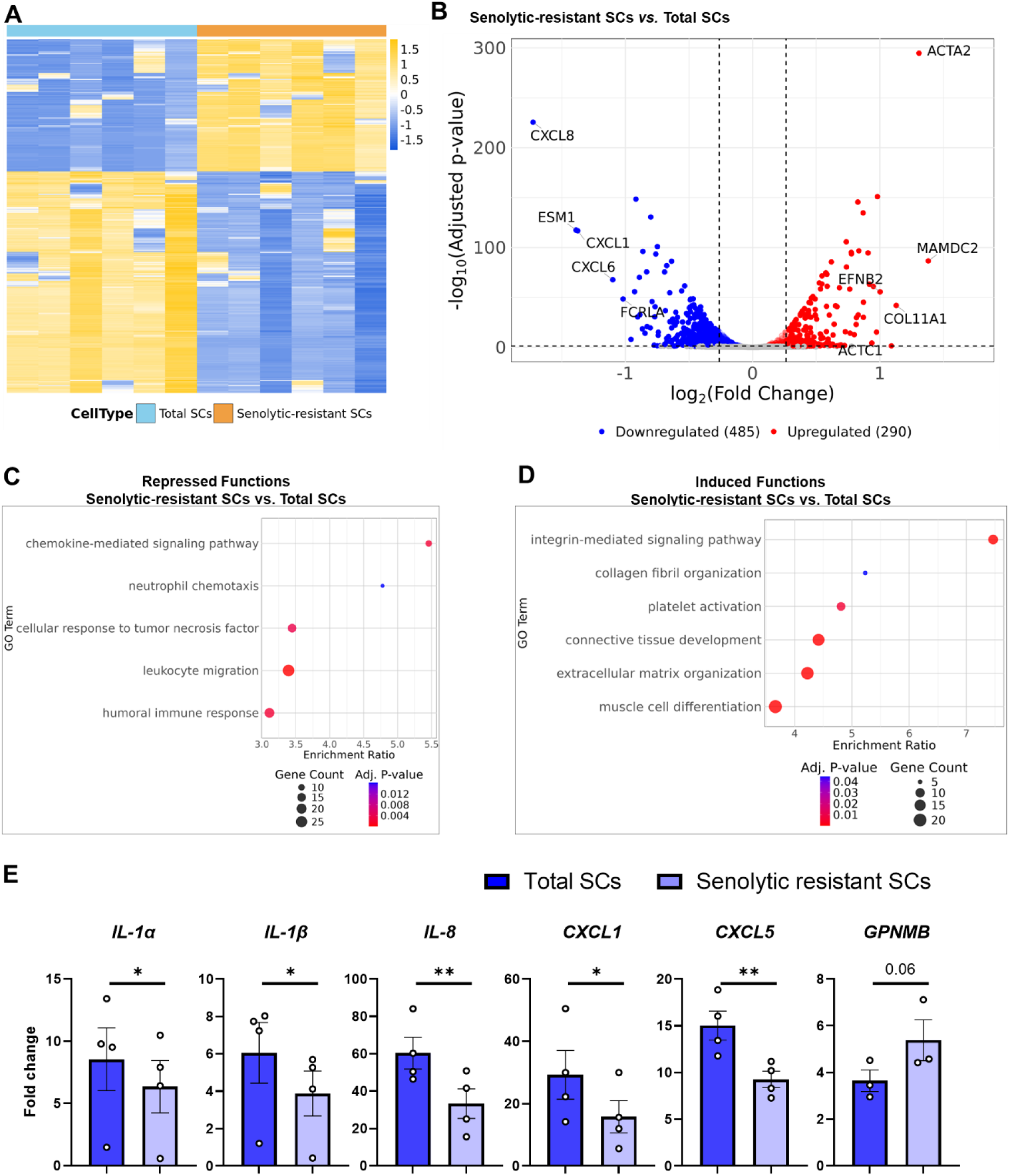
Senolytic-resistant human senescent preadipocyte SASP profiles are distinct from the total SC population. (A) Heat map of differentially expressed genes (DEGs). (B) Volcano plot of the DEGs indicating key genes. (C, D) Gene ontology analysis of biological functions of DEGs in the total *vs.* senolytic-resistant SC populations. (E) SASP factor expression (rt-PCR) in senescent preadipocytes. Data are expressed as a function of vehicle-treated non-senescent cells. Means +/- SEM; paired, 2-tailed Student’s T-tests.

Analogously to preadipocytes, senescent HUVECs resistant to Quercetin had differences in gene expression compared to the total HUVEC SC population, with a less inflamed gene expression pattern in the senolytic-resistant HUVECs than the total senescent HUVEC population (Fig. S6). With the sample size used in our studies, unlike SCs, non-senescent control human preadipocyte or HUVEC cultures exposed to senolytics did not have statistically significant reductions in their already low pro-inflammatory SASP factor expression compared to cells exposed to vehicle (Figs. S7A, B). Although some SASP factors were upregulated in non-senescent cells treated with senolytics, the levels of these pro-inflammatory factors remained lower than SCs. This observation suggests that while senolytics clear SCs and reduce inflammation over the long term, they can also elicit stress adaptation responses by short-term, transient activation of pro-inflammatory signals in non-senescent cells. These findings support the possibility that senolytics primarily target the SC subtype with an inflammatory, apoptosis-inducing SASP profile, consistent with the hypothesis-driven approach that we used to discover the first senolytics [29].

### Senolytic-Resistant Human Senescent Preadipocytes Differ in Extent of Induction of Inflammation and mt-DNA Secretion from the Total SC Population

Senescent preadipocytes can spread an inflammatory state to non-senescent cells and activate immune cells, in part through SC mtDNA production and release [41, 55]. This induction of inflammation by SC mtDNA was recently confirmed [31, 56]. Given the differences in SASP profiles between senolytic-resistant and the total SC populations, whether the SC subtypes induce inflammation to the same extent in non-senescent cells was tested (Fig. 5A). Non-senescent preadipocytes from healthy donors had lower expression of inflammatory factors when exposed to CM from senolytic-resistant cells compared to those cultured with CM prepared from the total SC population (Fig. 5B). Furthermore, cell-free mtDNA content in CM, marked by expression of NADH dehydrogenase subunit 2 (MT-ND2) and cytochrome c oxidase subunit III (MT-COX3) of the electron transport chain (known to be pro-apoptotic [56, 57]) tended to be higher in cultures of the total SC population than in cultures of the senolytic-resistant SC population (Fig. 5C), unlike cell-free nuclear DNA (Fig. S8). Hence, senolytic-resistant SCs appear to provoke less inflammation than the total population of SCs (senolytic-resistant *plus* senolytic-sensitive SCs).

**Fig. 5.**
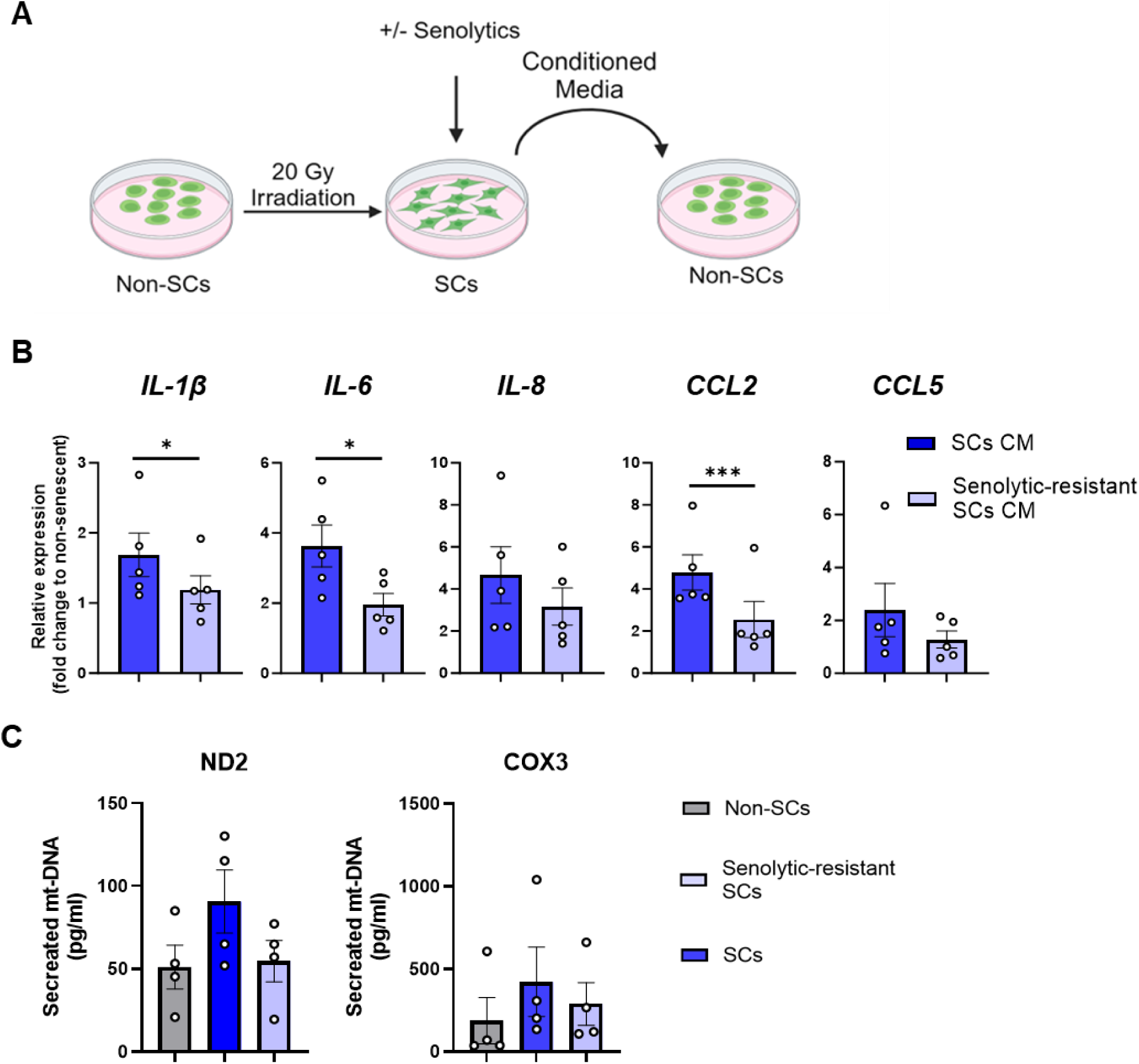
Effects of the senolytic-sensitive *vs*. total SC populations on induction of inflammation and secretion of mt-DNA. (A) Experimental scheme. (B) Non-senescent preadipocytes were treated with conditioned media (CM) from resistant *vs.* total SC populations for 24 hrs. and inflammatory factors were analyzed by rt-PCR. Means +/- SEM; paired, two-tailed Student’s T-tests. (C) Secreted mt-DNA by indicated cell types. Means +/- SEM; paired, 1-way ANOVA; *post hoc* pairwise comparison by Tukey’s HSD multiple comparison test.

### Transplanting Senolytic-Resistant Subtype SCs Causes Less Physical Dysfunction Than Transplanting the Total SC Population

Transplanting small numbers of SCs into mice is sufficient to induce features of frailty, suggesting a role of SCs in causing physical dysfunction [22]. To test whether the SC subtypes differ in their impact on physical function *in vivo*, 1×10^6^ senolytic-resistant or total population SCs were transplanted intraperitoneally into 2-month-old male young *SCID-Beige* mice (Fig. 6A). After 1-month, physical function variables were tested. In previously healthy mice that were transplanted with total SC population cells, reductions in grip strength (Fig. 6B) and hanging endurance (Fig. 6C) were more evident than in mice transplanted with senolytic-resistant SCs, indicating differences between the impacts of the SC subtypes on function.

**Fig. 6.**
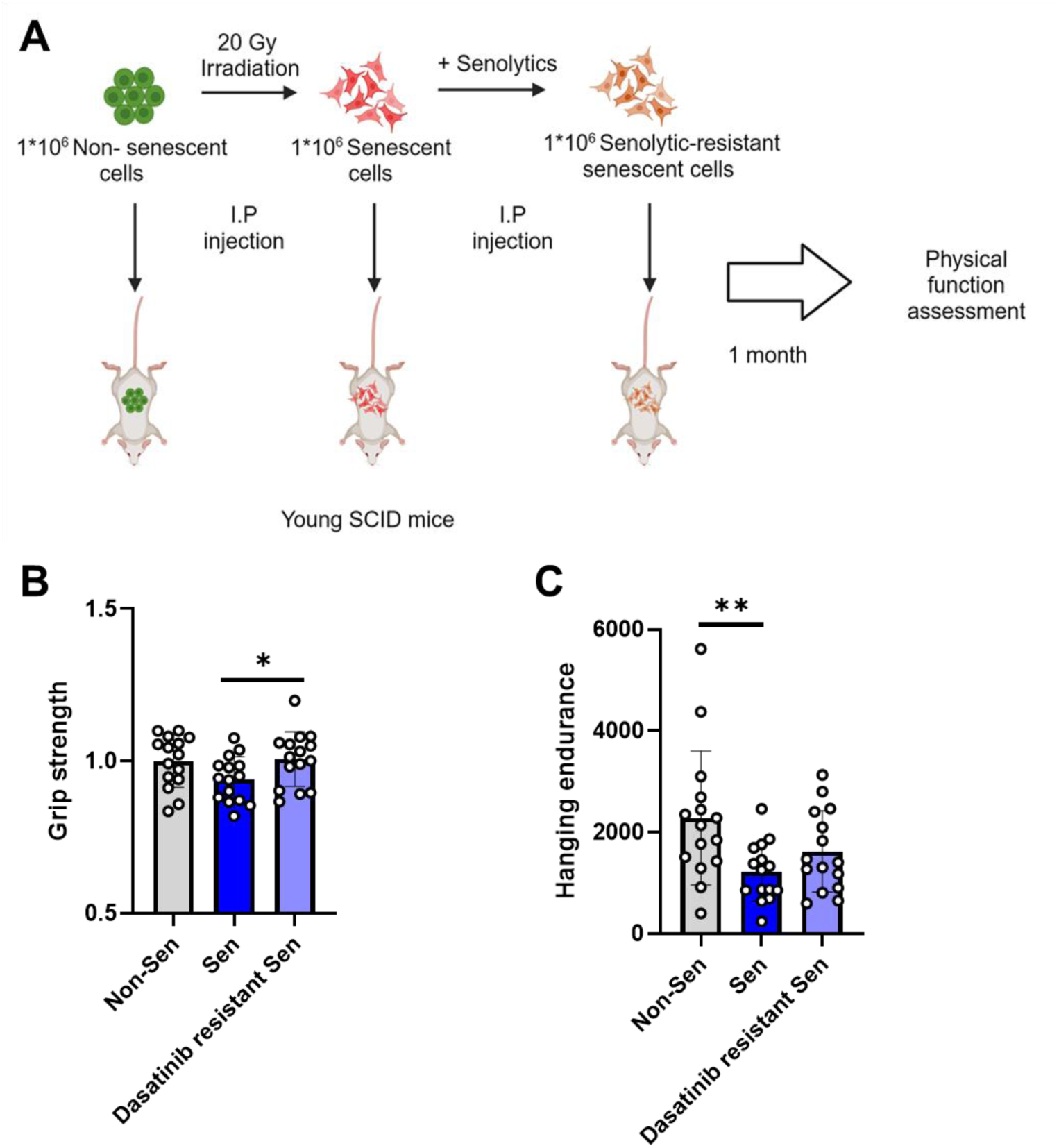
SC subtypes transplanted into younger mice differ in impact on physical function. (A) Experimental scheme. (B) Grip strength and (C) wire hanging endurance in 2-month-old male *SCID-Beige* mice 1 month after being transplanted with 1×10^6^ senolytic-resistant or total senescent human preadipocyte population cells by intraperitoneal injection (N=15). Baseline grip strength was similar in each group of mice before transplantation (Fig. S9). Data are shown as means +/- SEM with individual values; unpaired two-tailed Student’s T-tests.

### A “1:2:3:4”-Step Approach Enhances Ablation of Cultured Triple-Negative Breast Cancer Cells

Lastly, we explored the potential application of senosensitizers in the context of cancer. Triple-negative breast cancer (TNBC) remains one of the most challenging malignancies to treat due to its poor prognosis and resistance to conventional therapies. It has been found that chemotherapy can induce a senescent phenotype in cancer cells and that senolytics can eliminate some of these therapy-induced senescent (TIS) cells [58]. Based on these points and our findings above about senolytic-sensitive *vs*. -resistant SCs, we hypothesized senosensitizers could enhance effectiveness of senolytics in removing TIS. We previously found TLR3 agonists (which activate the PAMP pathway also activated by coronaviruses) upregulate pro-inflammatory/ pro-apoptotic SASP factors in senescent cells [59]. Therefore, as a preliminary test of our hypothesis, we treated human TNBC MDA-MB-231 cells with Cisplatin, followed by Dasatinib, then the TLR3 agonist polyinosinic-polycytidylic as a senosensitizer, and finally Dasatinib again (Fig. S10). This “1:2:3:4” stepwise regimen (chemotherapy → senolytic → senosensitizer → senolytic) resulted in a ∼90% further reduction in TNBC cell survival than cells treated with chemotherapy alone (Fig. S10).

## Discussion

Based on differences in SASP profiles, responsiveness to senolytics, and functional outcomes, we propose a conceptual framework distinguishing senolytic-sensitive and senolytic-resistant senescent cell subtypes. While these categories are not absolute or mutually exclusive, they provide a useful lens for interpreting observed heterogeneity in senescent cell behavior. Both have characteristics of SCs (increased p16^INK4a^, p21^CIP1^, γH2AX, SAβgal, and proliferative arrest), but the subtypes differ in SASP factors produced, mtDNA release, responsiveness to senolytics, and functional effects *in vitro* and *in vivo*. These “senolytic-sensitive” and “senolytic-resistant” SC subtypes are reminiscent of the “deleterious” and “helper” SC subtypes previously proposed [24]. The subpopulations might arise in response to pre-existing diversity within non-senescent cells, with effects being amplified once cells have become senescent through such mechanisms as: 1) expression of genes that had been silent in non-senescent cells due to epigenetic changes and associated chromatin openness in SCs, transpositional events, or other SC-intrinsic mechanisms, 2) differences in mtDNA abundance or localization between the senolytic-resistant and -sensitive SC subtypes (mtDNA is linked to inflammation in SCs [55]), 3) host *milieu*, including characteristics of cells near the SCs [60], or 4) extent of replication of cells before they originally became senescent, among other potential mechanisms or combinations of mechanisms. The senolytic-resistant SC subtype appears to have a more strongly pro-repair, fibrotic, and growth promoting profile, possibly in part related to TGF-β [61], than senolytic-sensitive SCs. The senolytic-sensitive subtype has a SASP that is inflammatory, with higher expression of cytokines and chemokines and more mtDNA release than the senolytic-resistant subtype.

The first senolytics were discovered based on the hypothesis that SCs resist apoptotic stimuli related to increased pro-survival, anti-apoptotic defenses and the finding that targeting these SC anti-apoptotic pathways leads to elimination of 30-70% of SCs [29]. Consistent with this, the SASP of senolytic-resistant SCs is less pro-inflammatory, pro-apoptotic, and tissue destructive than the SASP of senolytic-sensitive SCs. Additionally, suppressing or amplifying pro-inflammatory SASP features, respectively, decreased or enhanced extent of SC killing by senolytics. The interplay between the SASP and SCAPs and verification that Ruxolitinib or LPS modulated apoptosis (caspase assays, TUNEL) need to be explored in further studies. Single-cell proteomic studies in mice with obesity, a state marked by increased pro-inflammatory SC abundance especially in epididymal fat depots, indicated that within the senescent preadipocyte population (marked by p16+, p21^Cip1^+, γH2AX+), a cluster highly expressing pro-inflammatory/ pro-apoptotic factors was more effectively eliminated by senolytics than other cell populations. Furthermore, SCs can amplify inflammatory factor production in non-senescent cells [41] and here we found that it is principally the subset of senolytic-sensitive SCs that does so.

Infiltration of immune cells into adipose tissue together with increased SC abundance is a feature of both aging and obesity [41]. Senolytic treatment of obese mice led to reduced adipose tissue natural killer (NK) cells, which are an innate immune cell type that can eliminates persisting SCs [62]. This is reminiscent of the observation that migration of tail vein-injected labelled monocytes into adipose tissue of obese mice treated with senolytics is less than that in obese mice treated with vehicle [49]. Hence, senolytic-induced decreases in the burden of senolytic-sensitive SCs that produce inflammatory factors and chemokines may reduce immune cell infiltration and activation *in vivo*. This may occur despite the continued presence of senolytic-resistant SCs because these cells are less pro-inflammatory and may express more GPMNB than non-senescent cells, possibilities that merit further investigation.

We investigated the Dasatinib- and Quercetin-resistant *vs*. -sensitive human preadipocyte and endothelial SC subtypes, respectively. Potentially, other cell types becoming senescent or other senolytics could result in senolytic-resistant SC subsets with characteristics fundamentally distinct from those studied here. However, in keeping with findings in these two human cell types, a recent study indicated that a subset of senescent fibroblasts with a pro-inflammatory SASP profile related to NF-κB and the YAP-TEAD complex is more susceptible to the senolytic, VPF, than other senescent fibroblast subsets [63]. Further studies are needed to ascertain whether the senolytic-sensitive *vs*. -resistant SC subtypes with their apoptotic, tissue-damaging *vs*. pro-growth profiles, respectively, occur across more cell types and over a range of different senolytics.

Senolytic-resistant SCs could have detrimental effects. PAMPs generated during infections may rapidly alter the SASP of pre-existing senolytic-resistant SCs, with substantially increased release of tissue-damaging, inflammatory, pro-apoptotic factors, potentially worsening symptoms and contributing to complications, perhaps including cytokine storm. Indeed, LPS can amplify inflammatory SASP factors in SCs by over an order of magnitude within 3 hrs.; hence, effects of pathogenic infections on senolytic-resistant cells need to be examined [50]. In malignancies, therapy-induced senolytic-resistant SCs may produce growth factors that promote cancer growth, express immune evasion signals such as GPNMB, and release pro-fibrotic factors that create a matrix shielding cancers from immune clearance, speculations that also need to be explored. Senolytic-resistant cancer therapy (radiation or chemotherapy)-induced SCs (TIS) harboring cancerous mutations could lead to tumor relapse once these SCs have accumulated further mutations and escape senescence [15]. Therefore, there may be situations in which strategies for ablating senolytic-resistant “helper” SCs may be important over and above eliminating only pro-apoptotic/ pro-inflammatory SCs (Fig. S11).

Studies are needed to test whether these hypotheses are correct, because an asymptomatic reservoir of “helper”, senolytic-resistant SCs could build up with aging or after such senescence-inducing insults as trauma, obesity/ diabetes, chemotherapy, radiation, or infections, among others. To this end, because pro-apoptotic, inflammatory SASP features can be attenuated by senomorphics (*e.g*., Ruxolitinib [Fig. 1], Metformin [64], or Rapamycin [65]) and be rapidly upregulated by PAMPs (*e.g*., LPS or TLR3 activation in response to coronavirus [50] [59]), we speculate that the impact of senolytics might be reduced by senomorphics and enhanced by agents related to PAMPs (Fig. S11). If senomorphics do indeed impede effectiveness of senolytics, this would imply that simultaneous (as opposed to sequential) administration of senolytics and senomorphics may have less than additive effects in delaying, preventing, alleviating, or treating senescence-related disorders and diseases. Conversely, accentuating apoptotic SASP factor production by exposing senolytic-resistant SCs to agents currently under investigation, “senosensitizers”, which act through PAMP-related mechanisms, may sensitize previously senolytic-resistant SCs to senolytics. Potentially, this might defend against complications of insults such as infections in individuals harboring previously clinically silent senolytic-resistant SCs. In patients with malignancies undergoing chemotherapy/ radiation, we also speculate that administration of senosensitizers could boost the numbers of cancer-harboring TIS that can be removed subsequently by senolytics (Fig. S10). In this scenario, effectiveness of standard chemotherapy/ radiation could possibly be augmented by the proposed “1-2-3-4 punch” approach: 1) Chemotherapy/ radiation; 2) A round of senolytics to remove senolytic-sensitive TIS; 3) A round of senosensitizers to enhance responsiveness to senolytics of any remaining cancer-harboring, previously senolytic-resistant TIS; and 4) An additional round of senolytics to remove these now senolytic-sensitive remaining cells. Early data supporting hypothesis in the case of triple negative breast cancer are shown in Fig. S10. Hence, there is a pressing need to comprehensively characterize senolytic-resistant SCs and develop strategies to eliminate these cells, including developing effective senosensitizers and therapeutic regimens entailing sequencing administration of senosensitizers and senolytics. Although the data presented in this manuscript are based on a limited number of subjects and cell types, which may raise concerns regarding generalizability, they have the potential to establish a new paradigm in cancer and senescence-targeted therapies. In our view, these provocative findings warrant further investigation.

## Author Contributions

U.T., T.T., and J.L.K. generated the overall concept of the study. U.T., M.S., C.I., A.P, V.K., K.J., T.P., S.C., L.P., and Y.Z. performed experiments. U.T, M.S., T.T., and J.L.K. designed the study and wrote the manuscript. B.T.P., H.K.Y., and R.M. analyzed the RNA-seq data. N.G. and M.X. assisted with RNA-seq. N.G. conducted the isolation of preadipocytes from human fat tissues. D.B.A., H.K.Y., and R.M. supervised the statistical analyses. All authors read, edited, and approved the final version of the manuscript.

## Acknowledgements

The authors are grateful to the subjects who allowed us to harvest adipose tissue at the time of their kidney transplant donation surgery or undergoing gastric bypass surgery and to Mrs. Marcia K. Mahlman for assistance in acquiring these samples from which human preadipocytes were isolated. We are grateful to bioinformatics core at the University of Connecticut for assistance with primary and secondary analysis of RNA-sequencing data.

## Data Availability

All raw data used to generate graphs are included in Supplemental Data. Raw sequencing data and gene counts are available through NCBI’s Gene Expression repository under accession number: GSE268701.

---

**Fig. S1.**
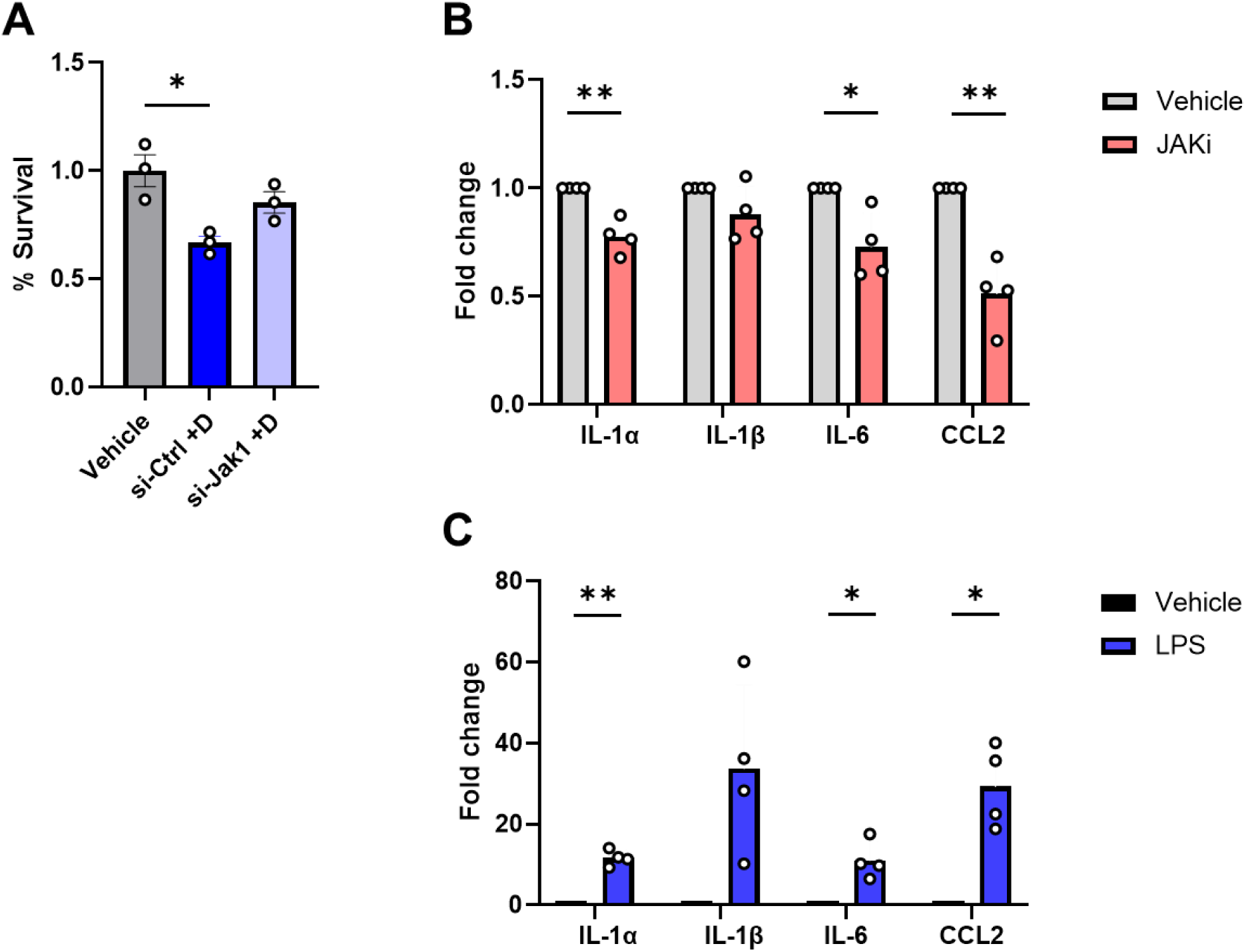
Effect of SASP modulation on senescent cell survival and pro-inflammatory factors. (A) % Survival of senescent preadipocytes. Means +/- SEM, 1-way ANOVA; *post hoc* comparisons by Tukey’s HSD multiple comparison test. (B) JAKi and (C) LPS modulate SASP expression (rt-PCR) of senescent preadipocytes. Data are expressed as a function of vehicle-treated cells. Means +/- SEM, paired, two-tailed Student’s T-tests.

**Fig. S2.**
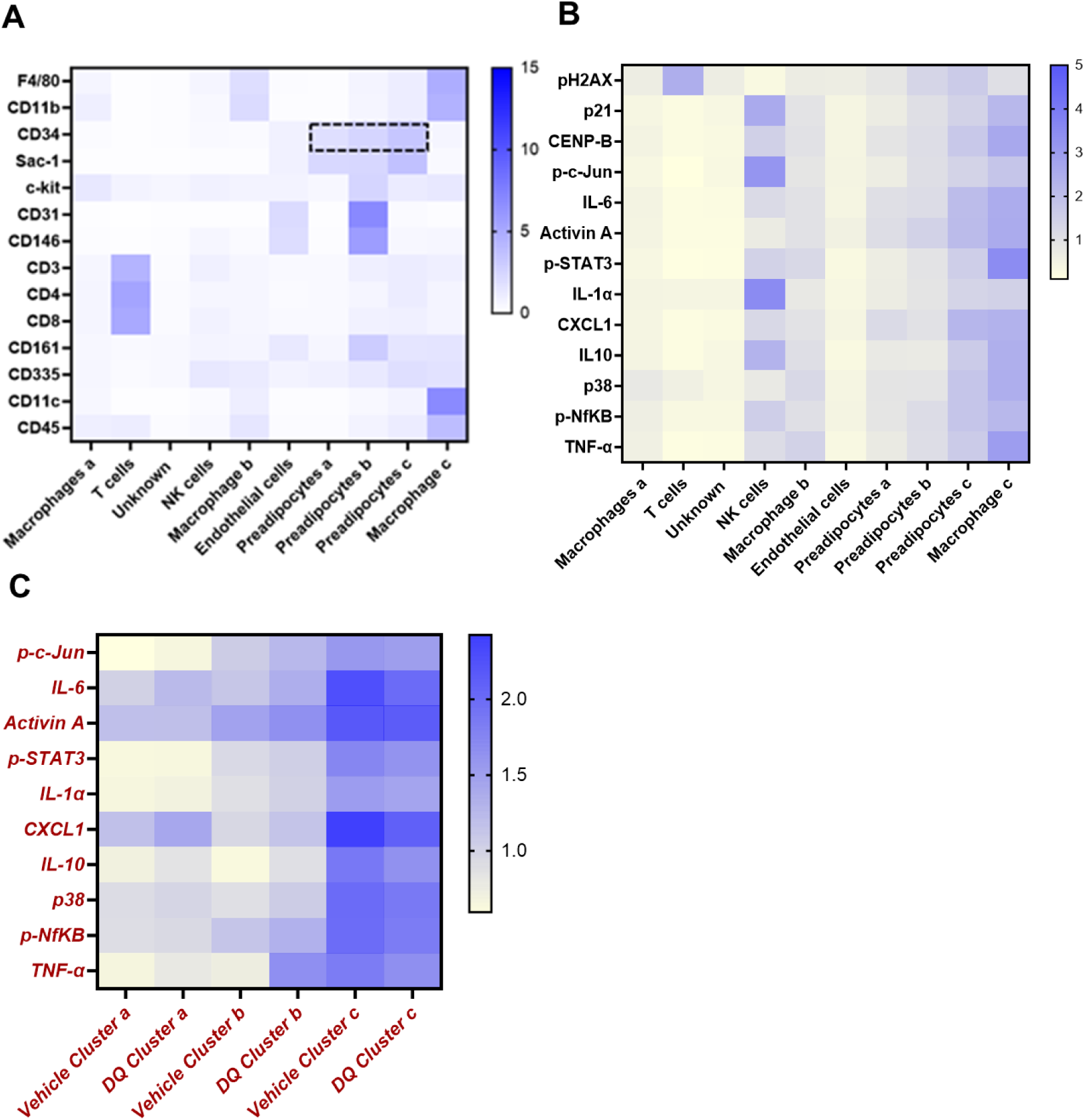
Heatmap of the 10 cell clusters in Fig. 2B with cell identification markers (A) and SASP proteins (B). The preadipocyte cluster is highlighted. (C) SASP profiles in vehicle *vs*. Dasatinib plus Quercetin-treated preadipocytes.

**Fig. S3.**
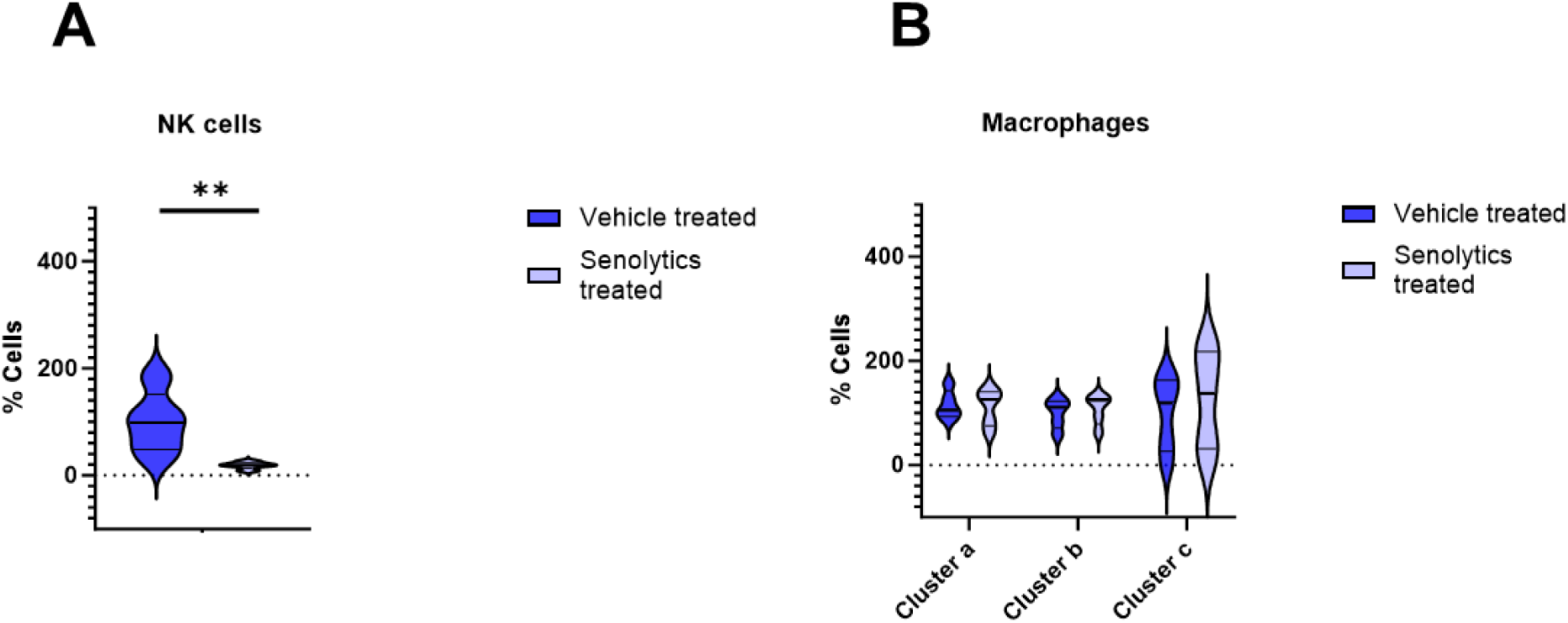
Senolytic treatment alters NK cell adipose tissue content. (A) Percent of NK cells in each cluster in vehicle-*vs*. senolytic-treated obese mice. (B) Percent of macrophages in vehicle-*vs*. senolytic-treated obese mice. Means +/- SEM; unpaired two-tailed Mann-Whitney tests.

**Fig. S4.**
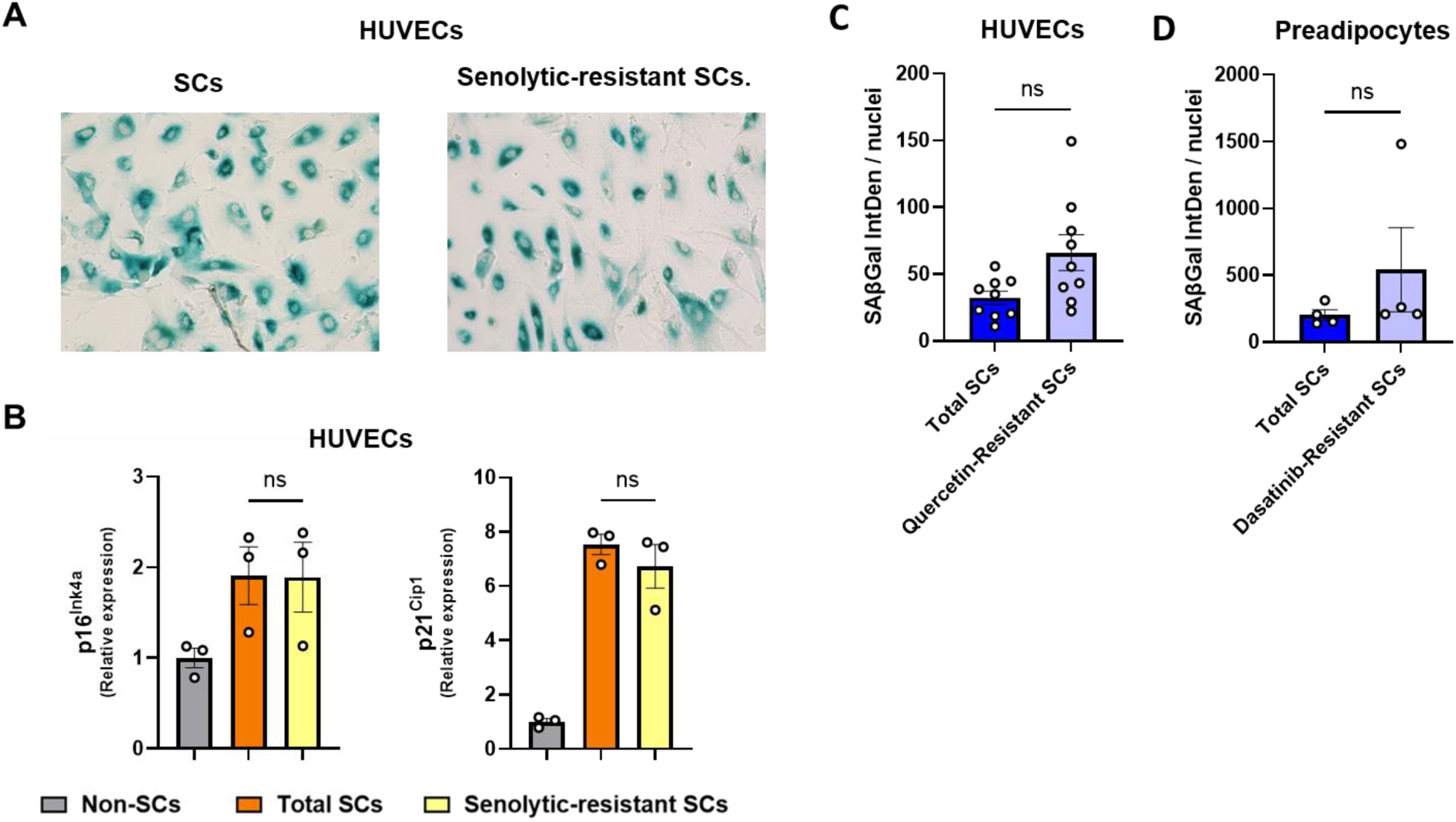
Supplemental data related to Fig. 3. As in human senescent preadipocytes, senescence markers are similar in the human endothelial cell (HUVEC) senolytic-resistant *vs*. total SC populations. Human non-senescent or senescent HUVECS were incubated with vehicle or Quercetin 10 µM for 24 hrs., washed 4X, and maintained in senolytic-free medium for a further 72 hrs. (A) SAβgal was analyzed. (B) Expression of the indicated senescence markers (N=3) in the total HUVEC SC population *vs*. senolytic-resistant senescent HUVECs. Data are expressed as a function of non-senescent control cells; means +/- SEM; unpaired, 1-way ANOVA; *post hoc* comparisons by Tukey’s HSD multiple comparison test. (C) Quantification of SAβgal intensity in senescent HUVECs and (D) preadipocyte subtypes. Means +/- SEM; paired two-tailed Student’s T-tests.

**Fig. S5.**
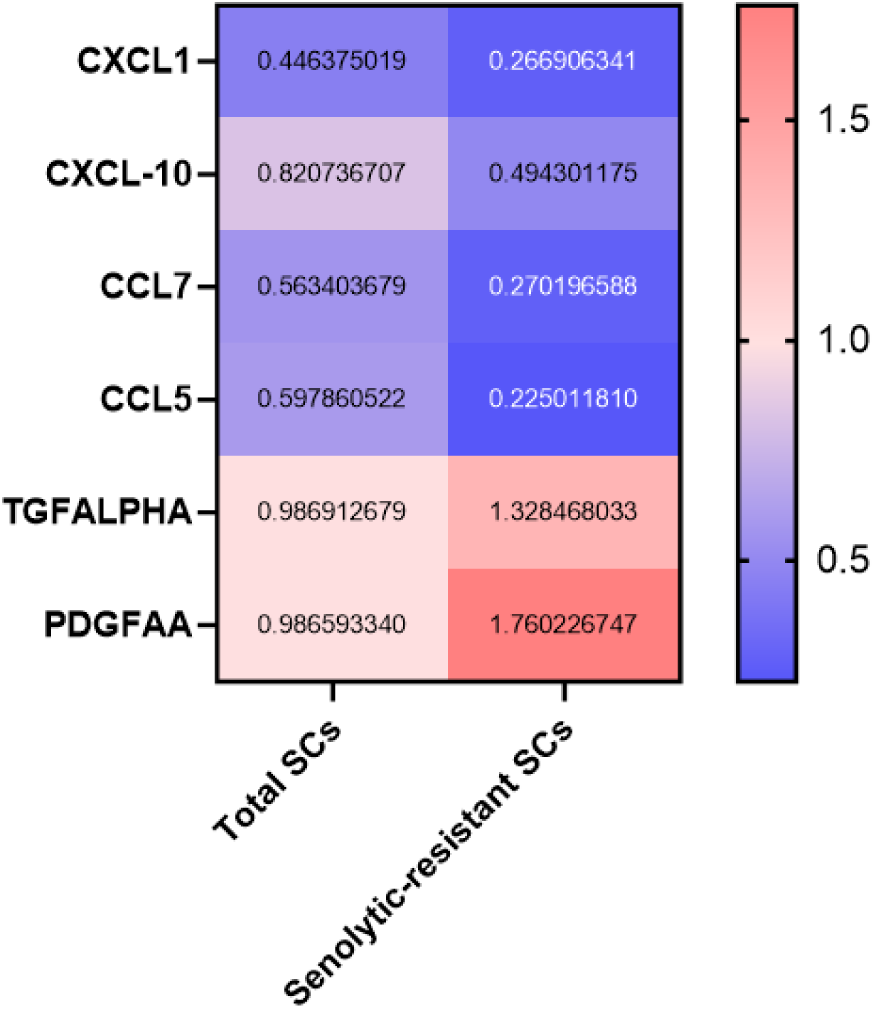
Heat map of SASP factor expression in conditioned media collected from the total senescent cell or senolytic-resistant senescent cell preadipocyte populations (n=3). Cells were treated with Dasatinib for 3 days. Multiplex ELISA was performed in conditioned media collected after 72 hrs. after washing out Dasatinib. Relative abundance in senolytic-resistant *vs*. total SCs; results normalized to the mean of vehicle-treated total SCs.

**Fig. S6.**
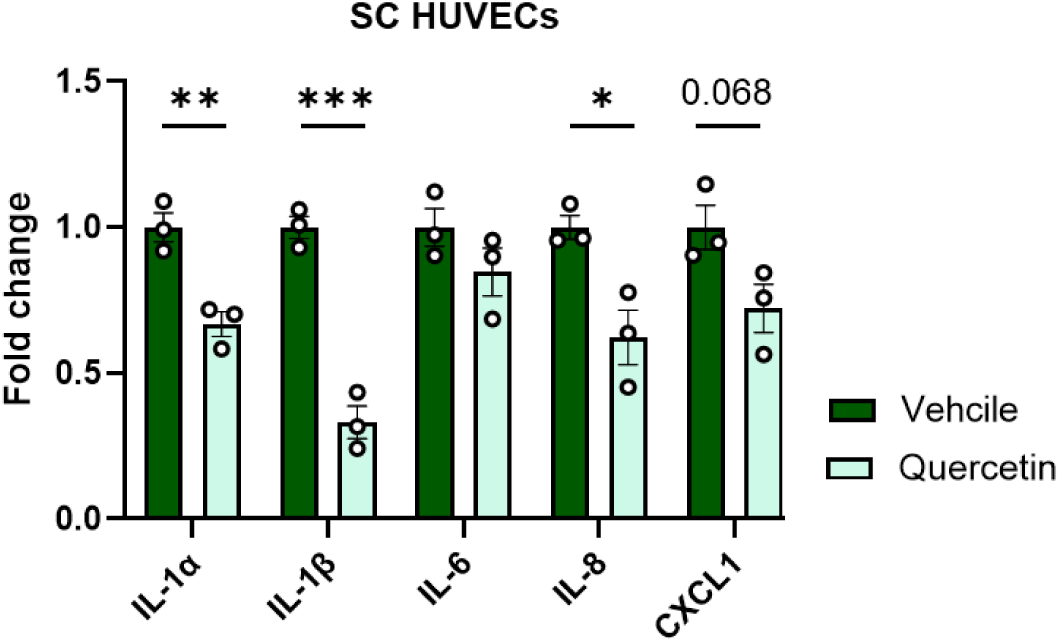
Quercetin-resistant senescent HUVECs have reduced SASP factor expression (rt-PCR). Data are expressed as a function of vehicle-treated cells. Means +/- SEM; unpaired two-tailed Student’s T-tests.

**Fig. S7.**
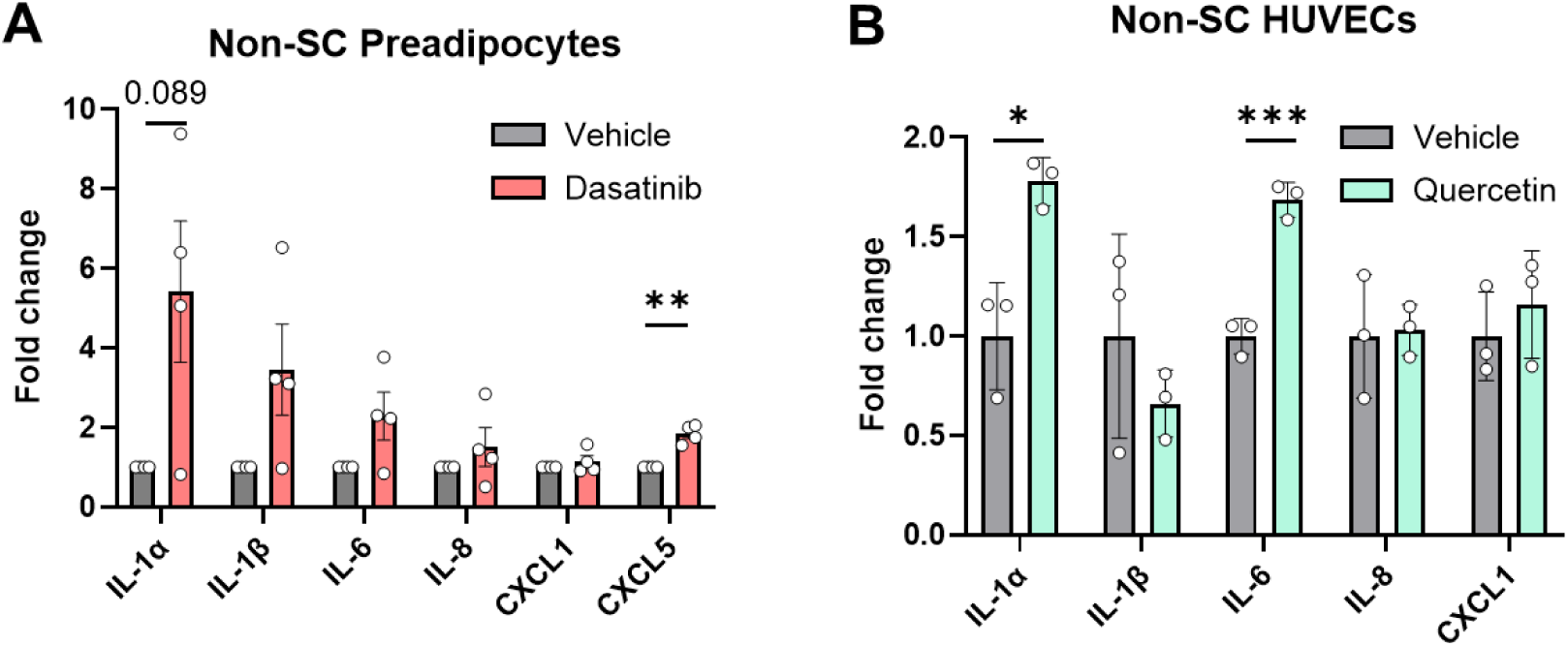
Senolytic effects on pro-inflammatory factors in non-senescent preadipocytes and HUVECs. (A) Non-senescent human preadipocytes treated with Dasatinib expressed as a function of preadipocytes treated with vehicle. (B) Non-senescent HUVECs treated with Quercetin expressed as a function of HUVECs treated with vehicle. Data are expressed as a function of vehicle-treated cells; means +/- SEM; paired (A) or unpaired (B), two-tailed Student’s T-tests.

**Fig. S8.**
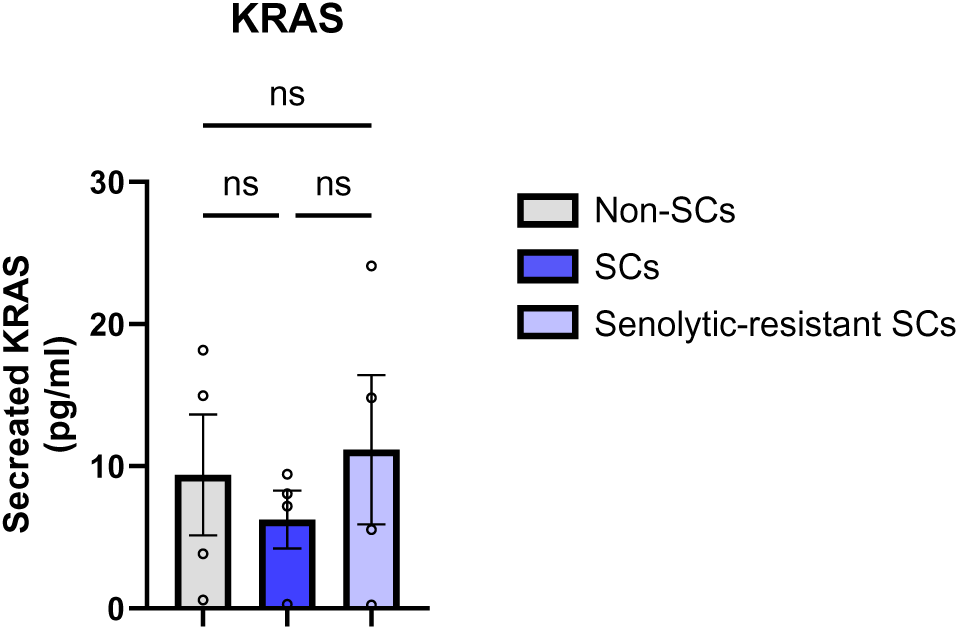
Cell-free nuclear DNA release by the non-senescent *vs*. the total senescent (senolytic-resistant <u>plus</u> senolytic-sensitive) *vs*. senolytic-resistant human preadipocyte populations. Abundance of cell-free KRAS DNA in CM prepared from the indicated cell types. Means +/- SEM. No statistically significant differences (ns) were detected among three groups by paired, 1-way ANOVA with *post hoc* pairwise comparisons by Tukey’s HSD multiple comparison test.

**Fig. S9:**
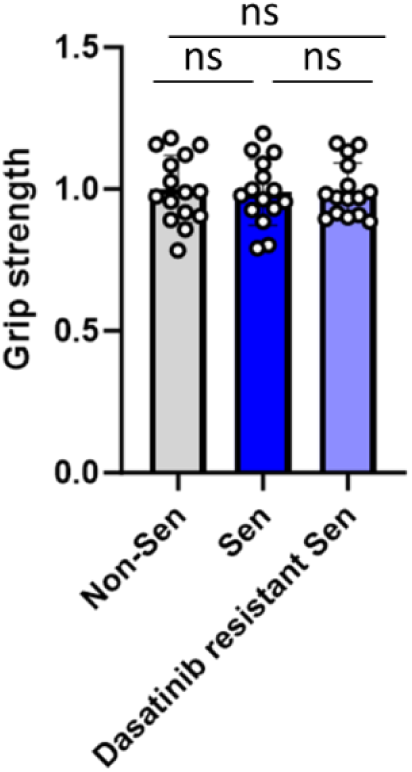
Baseline grip strength prior to transplantation of cells. Base line grip strength in 2-month-old male *SCID-Beige* mice (N=15). Data are shown as means +/- SEM with individual value. No statistically significant differences (ns) among three groups by unpaired two-tailed Student’s T-tests.

**Fig. S10:**
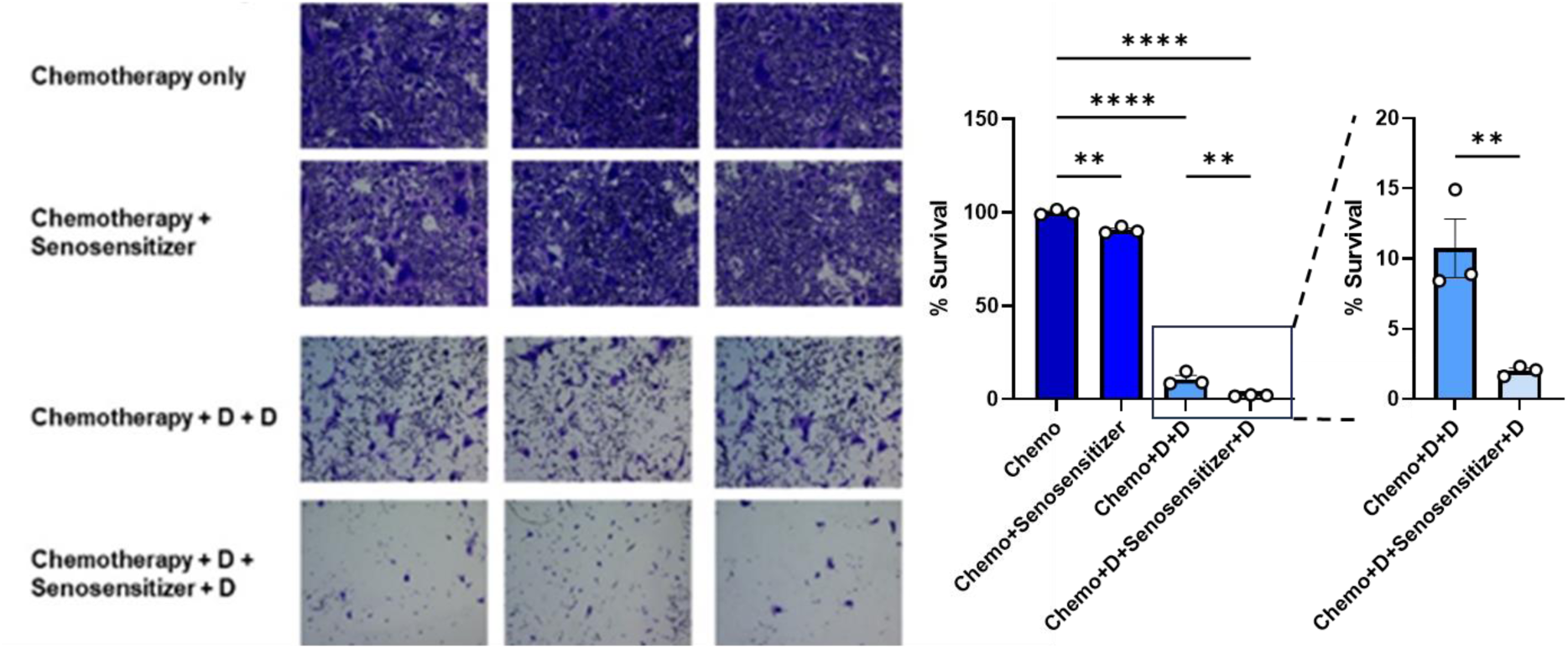
A “1:2:3:4”-step approach enhances ablation of cultured TNBC cells. Representative images of surviving (crystal violet+) MDA-MB-231 human breast cancer cells and quantification relative to chemotherapy only-treated cells. Chemo: Cisplatin (1µ/ml), Senosensitizer: TLR3 agonist (polyinosine-polycytidylic acid) (10 µg/ml), D: Dasatinib (200 nM). This “1:2:3:4” step regimen was repeated 3 times. These findings provide early evidence that a sequential therapeutic approach incorporating chemotherapy, senosensitizers, and senolytics could offer a promising strategy to overcome resistance in aggressive cancers such as TNBC, a possibility that merits further study. Data are shown as means +/- SEM with individual values; 1-way ANOVA; *post hoc* pairwise comparisons by Tukey’s HSD multiple comparison test.

**Fig. S11.**
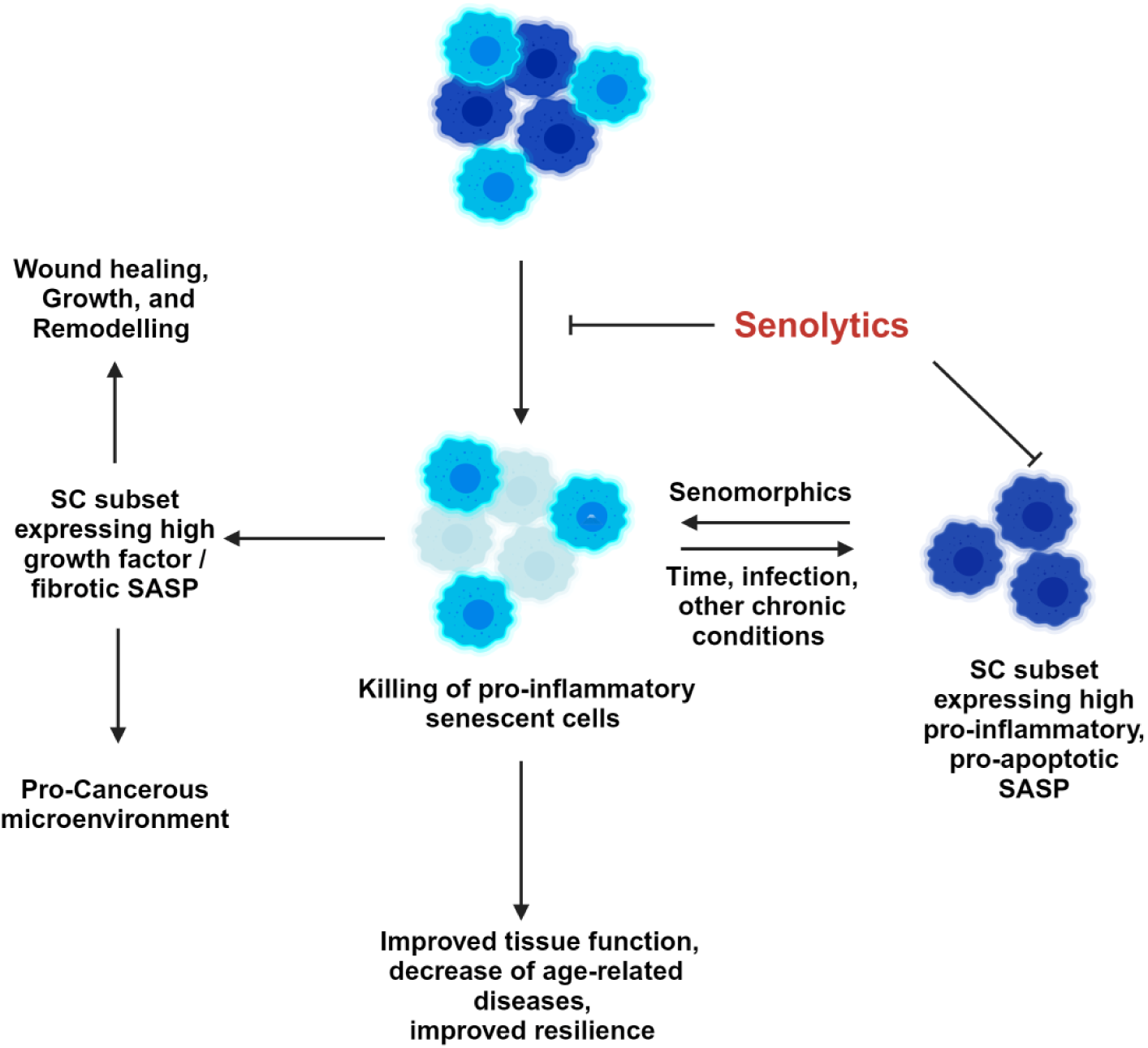
Summary and implications. The SC subtype with a pro-inflammatory, pro-apoptotic, tissue-damaging SASP is susceptible to senolytics. The SC subset not killed by senolytics have a pro-growth/ repair SASP with less mtDNA release and less proclivity to cause physical dysfunction following transplantation into younger individuals than the total SC population. On the one hand, agents such as Ruxolitinib can attenuate the pro-inflammatory SASP, making SCs become resistant to senolytics. On the other, insults such as infections (exposure to PAMPs) can accentuate inflammatory SASP features within hours [50], at least *in vitro*. Therefore, we propose it may be feasible to develop agents, “senosensitizers”, that induce senolytic-resistant SCs to be converted into SCs that are susceptible to senolytics. Sequencing senosensitizers with senolytics could enable removal of a pool of relatively silent SCs harbored in apparently asymptomatic individuals, which could otherwise be activated within hours by infections or other insults into deleterious cells.

